# Constitutively synergistic multiagent drug formulations targeting MERTK, FLT3, and BCL-2 for treatment of AML

**DOI:** 10.1101/2023.03.13.531236

**Authors:** James M Kelvin, Juhi Jain, Aashis Thapa, Min Qui, Lacey A Birnbaum, Samuel G Moore, Henry Zecca, Ryan J Summers, Emma Costanza, Biaggio Uricoli, Xiaodong Wang, Nathan T Jui, Haian Fu, Yuhong Du, Deborah DeRyckere, Douglas K Graham, Erik C Dreaden

**Author notes:** corresponding authors Correspondence to (ECD) and (DKG). equal contribution (James M Kelvin, Juhi Jain). Department of Pediatrics, University of Arizona College of Medicine, and Banner University Medical Center Tucson, Tucson, AZ 85724, USA.

## Abstract

Although high-dose, multi-agent chemotherapy has improved leukemia survival rates in recent years, treatment outcomes remain poor in high-risk subsets, including acute myeloid leukemia (AML) and acute lymphoblastic leukemia (ALL) in infants. Development of new, more effective therapies for these patients is therefore an urgent, unmet clinical need. To address this challenge, we developed a nanoscale combination drug formulation that exploits ectopic expression of MERTK tyrosine kinase and dependency on BCL-2 family proteins for leukemia cell survival in pediatric AML and *MLL-*rearranged precursor B-cell ALL (infant ALL). In a novel, high-throughput combination drug screen, the MERTK/FLT3 inhibitor MRX-2843 synergized with venetoclax and other BCL-2 family protein inhibitors to reduce AML cell density *in vitro*. Neural network models based on drug exposure and target gene expression were used to identify a classifier predictive of drug synergy in AML. To maximize the therapeutic potential of these findings, we developed a combination monovalent liposomal drug formulation that maintains ratiometric drug synergy in cell-free assays and following intracellular delivery. The translational potential of these nanoscale drug formulations was confirmed in a genotypically diverse set of primary AML patient samples and both the magnitude and frequency of synergistic responses were not only maintained but were improved following drug formulation. Together, these findings demonstrate a systematic, generalizable approach to combination drug screening, formulation, and development that maximizes therapeutic potential, was effectively applied to develop a novel nanoscale combination therapy for treatment of AML, and could be extended to other drug combinations or diseases in the future.

## INTRODUCTION

Acute leukemia is a hematopoietic malignancy characterized by uncontrolled proliferation of undifferentiated lymphoid precursors (ALL) or clonally heterogeneous myeloid blasts (AML). Despite gradual improvements in outcomes for patients with these subtypes of leukemia, 5-year survival rates are only 65-70% for pediatric patients after diagnosis with AML and less than 60% for infants (<1 year of age) with ALL (1, 2). Development of new, more effective therapies for AML and infant ALL is thus an urgent and unmet clinical need.

To address this challenge, we and others have developed strategies to improve potency and/or specificity of conventional drug modalities (3-13). Anthracyclines, for example, have been formulated in lipid nanoparticles (NPs) and have improved complete response rates (CRR) when combined with multiagent chemotherapy (69% CRR) in pediatric patients with relapsed AML compared to multiagent chemotherapy alone (59% CRR) (14). Fixed ratio liposomal formulations of cytarabine and daunorubicin (i.e. Vyxeos) have also been developed to prolong synergistic drug interactions and provided CRs in 54% of pediatric patients with relapsed or refractory (R/R) AML, with an 81% overall response rate and ∼97% of responders successfully transitioning to hematopoietic stem cell transplant (15).

Molecularly targeted inhibitors of the anti-apoptotic BCL2-family proteins have also been utilized to increase treatment efficacy and specificity. The BCL-2-selective inhibitor venetoclax prolonged survival in a murine xenograft model of *MLL*-rearranged (KMT2A) ALL, a mutation that occurs in 80% of infant ALLs and is associated with chemoresistance, increased risk of relapse and poor outcomes (16-18). Targeted modulation of the apoptotic pathway is also a strategy with particular relevance to AML due to its dysregulation during chemotherapy resistance (19). For instance, venetoclax has monotherapy activity in patients with R/R AML (20) and is FDA approved in combination with azacitidine, decitabine, or low-dose cytarabine for treatment of AML in older adults (21, 22). These favorable interactions with nucleoside analogs, chemotherapy, and hypomethylating agents suggest roles for BCL-2 in resistance to therapeutic agents with different mechanisms, or perhaps to cell stress in general, and demonstrate the utility of BCL-2 family inhibitors to sensitize leukemia cells to concurrently administered therapies (23, 24). Thus, venetoclax and/or other BCL-2 family inhibitors may synergize with a wide variety of other therapies to mediate leukemia cell killing.

Here, we investigate the potential of combined treatment with BCL2-family inhibitors and MRX-2843, a first-in-class small molecule inhibitor that targets two different leukemogenic tyrosine kinases (25, 26). One of these is FLT3, which is overexpressed in infant ALL (18) and activated by internal tandem duplication (FLT3-ITD) mutations (FLT3-ITD) in 15–25% of pediatric AMLs and 20–30% of adult AMLs (27, 28), where it is a validated therapeutic target, *e*.*g*. midostaurin and giltertinib are FDA approved for treatment of FLT3-mutant AML (29, 30). FLT3-ITD mutations have also been associated with poorer response rates to induction chemotherapy and inferior survival (28, 31) and they confer resistance to venetoclax by upregulating BCL-XL and MCL-1 (32, 33). MRX-2843 also targets MERTK, which is ectopically expressed in >80% of pediatric and adult AML and 30% of pediatric B-ALL patient samples (34-36). Transgenic expression of MERTK in the hematopoietic compartment leads to development of leukemia. Like the BCL2-family proteins, MERTK has a pro-survival role in leukemia cells, particularly in the context of cell stress, and mediates resistance to a variety of therapies (34, 37, 38). Pharmacologic inhibition of MERTK and/or FLT3 may synergize with BCL-2 family inhibitors to prevent AML progression. To investigate the therapeutic potential of combined MERTK/FLT3 and BCL-2 inhibition, we used a systematic approach to high-throughput combination drug screening and nanoscale drug formulation to maximize synergy between MRX-2843 and venetoclax and developed a transcriptomic classifier that predicts synergistic interactions and treatment response in cell culture assays. Both the frequency and magnitude of drug synergy were preserved in AML patient samples treated with the nanoscale drug formulation.

## MATERIALS AND METHODS

### Cell Lines and Cell Culture

KOPN-8 was provided by Dr. Christopher Porter (Emory University). RS4;11, KG1, THP-1 and MOLM-14 were provided by Dr. Lia Gore (University of Colorado Denver). SEM and NOMO-1 were from the Leibniz Institute *DSMZ*-German Collection of Microorganisms and Cell Cultures (Braunschweig, Germany). OCI-AML5, MV4-11, and Kasumi-1 were obtained from the American Type Culture Collection (Manassa,VA). All cell lines were cultured in RPMI 1640 with L-Glutamine (Corning; Corning, NY) supplemented with 10% heat-inactivated fetal bovine serum (Avantor Seradigm, VWR; Radnor, PA) and 1x Penicillin-Streptomycin (Corning) (cRPMI) except SEM cells were cultured in DMEM with L-Glutamine, glucose, and sodium pyruvate (Corning) instead of RPMI. All cells were cultured at 37°C in a 5% CO_2_ humidified atmosphere (Napco Series 8000 WJ CO_2_ Incubator; Thermo Electron Corporation, Waltham, MA) and tested at least every 4 months for mycoplasma. Live cells were counted (Countess III Automated Cell Counter, Invitrogen, Waltham, MA; Cellometer Auto T4, Nexcelom, Lawrence, MA) using Trypan blue (17-942E, Lonza; Conley, GA).

### High-throughput Ratiometric Drug Screening

MRX-2843 (Meryx, Inc.; Chapel Hill, NC), venetoclax (MedChemExpress; LC Laboratories, Woburn, MA), gossypol (AT-101; APExBIO, Taiwan) and obatoclax (LC Laboratories) were dissolved in dry DMSO (Sigma-Aldrich; Burlington, MA) and titrated in separate 384-well polypropylene plates (VWR) covered in sterile adhesive film (VWR) for storage prior to serial experiments. MRX-2843 drug arrays were plated orthogonal to those on BCL-2i plates such that high-throughput assembly of plate contents made both monotherapy dose series and checkerboard matrices comprising >150 unique pairwise ratios spanning 4 – 6 orders of magnitude. To screen cells against drugs, a Multidrop Combi Reagent Dispenser (Thermo Fisher; Waltham, MA) deposited 10,000 viable cells per well (0.25×10^6^/mL) in 384-well microplates (Greiner Bio-One; VWR). Cells were combined with drug matrices using a Beckman NX Liquid Handler (Beckman Coulter; Brea, CA) at the Emory Chemical Biology Discovery Center. Each well contained DMSO vehicle (Sigma-Aldrich) at 0.5% (v/v). Microplates were then centrifuged at 135 x g (Eppendorf 5810R; Eppendorf, Hamburg, Germany) for 5 minutes prior to incubating for 72 hours at 37°C in a 5% CO_2_ humidified atmosphere with gas permeable plate sealing film (VWR). Following incubation, cells were treated with the CellTiter-Glo 2.0 Viability Assay (1:2 v/v; Promega, Madison, WI) using the Viaflo Assist Pipetting Platform (Integra Biosciences; Hudson, NH). Cell microplates were mixed in a Microplate Mixer (Fisher) for 1 hour prior to measuring ATP-driven luminescence using a SpectraMax iD3 plate reader (Molecular Devices; San Jose, CA). HTS data quality was evaluated with the following acceptance criteria: CV<20% (positive and negative controls), SNR≥5 (positive controls), and Z’≥0.5 (per plate and cell line). Data was background-corrected to empty wells containing media/DMSO vehicle and normalized to positive controls, *i*.*e*. cells in media/DMSO vehicle. The impact of treatment on cell density (growth inhibition, GI) was computed as the complement of dose response viability. The Bliss Independence model was used to calculate growth inhibitory drug synergy. Additionally, we applied the Response Additivity model of drug synergy for predictive analyses. All drug combinations were tested in quadruplicate.

### Hierarchical Clustering

Hierarchical clustering was performed on cell line dose response data to select the pairwise combination that was maximally synergistic. Synergy scores were summed over the full response surface (SUS) of pairwise combinations within each cell line. SUS scores were grouped by agglomerative hierarchical clustering using Spearman rank distance (one minus spearman rank correlation) and average linkage between clusters with Morpheus matrix visualization software (Morpheus, https://software.broadinstitute.org/morpheus).

### Principal Component Analysis

PCA was performed on 7 variables of HTS-observed AML cell line responses. Combination variables of MRX-2843 and venetoclax consisted of total drug (Log_2_(M+V)) and drug ratio (Log_2_(M:V)), where M and V stand for concentrations (nM) of MRX-2843 and venetoclax, respectively. This data was combined with GI and synergy values (Bliss Independence) tabulated for each drug combination, along with publicly available log-transformed RNA expression data (Log_2_(TPM+1)) of MERTK, FLT3 and BCL-2 (Cancer Cell Line Encyclopedia, Expression 22Q2 Public; Broad Institute, Cambridge, MA) to distinguish cell lines. Data was standardized (mean=0, SD=1) to account for unequal SD across variables. We used Parallel Analysis with 1,000 Monte Carlo simulations of randomized data with equal dimension to set confidence intervals for PC eigenvalues. PCs attaining eigenvalues >95^th^ percentile of randomized confidence intervals were selected (GraphPad Prism version 9.5.0 for Windows, GraphPad Software, San Diego, California USA, www.graphpad.com).

### Neural Network Predictive Models

Variables used in PCA were implemented as predictor variables in a fully connected two-layer perceptron to train predictive models of continuous drug responses and classifiers of synergy. The NN consisted of two hidden layers comprising 3 TanH, 3 Linear and 3 Gaussian activation nodes each. 10 randomly generated split vectors partitioned the data into Training:Validation:Test sets with a 70:15:15 assignment, respectively. The neural network was allowed to tour the Training dataset over 100 times using the Squared Penalty fitting method at a fixed learning rate. For continuous variable prediction, a Robust Fit option was applied to eliminate the influence of outlier data, *i*.*e*. least squares were replaced by least absolute deviations to train the model. Synergy was calculated using the Bliss Independence and Response Additivity models. Synergy was rendered either as a continuous response variable or as a discrete category with the following classifications: observed (obs) responses ≥1% of expectation (exp) were labeled “Synergistic;” obs ≤ -1% exp were “Antagonistic;” all remaining responses were “Additive.” After training and testing predictive models, Monte Carlo simulations resampled predictor variables as independent inputs to assess their predictive value when alone and in combination with other predictor variables. Statistics of the predictor variable importance index, Total Effect, and overall model accuracy were compiled for each of the 10 split vector partitions (39). All neural network modeling and analyses, including ROC curve plotting, was done using JMP Pro, Version 16 (JMP©, Version 16, SAS Institute Inc., Cary, NC, 1989-2021).

### pH-Gradient Drug Loading into Lipid Nanoparticles

Liposomal NPs encapsulating MRX-2843 or venetoclax were made by first mixing organic solutions of DSPC (1,2-distearoyl-sn-glycero-3-phosphocoline; NOF Corporation, Shanghai, China), DSPG (1,2-distearoyl-sn-glycero-3-phosphoglycerol, sodium salt; NOF) and cholesterol (Sigma-Aldrich) lipids together at a 7:2:1 mol:mol ratio, respectively. DSPC and cholesterol were initially dissolved in chloroform (25 mg/mL), while DSPG (5 mg/mL) was dissolved in a chloroform:MeOH mixture (5:1, v/v). After mixing, solvent was removed under rotary evaporation and vacuum at 30°C (Rotavapor R-100; Buchi, New Castle, DE). Lipid foams were placed under vacuum overnight to remove residual solvent. Lipid foams were then rehydrated with acidic ammonium sulfate (EMD Millipore; Burlington, MA) solution (pH = 4.25, 500 – 600mM) and sonicated under high power via cup horn at 60°C (Q700, #431C2; Qsonica, Newton, CT). Resulting multilamellar vesicles were serially extruded under high pressure N_2_ gas (Liposofast LF-50; Avestin, Ottawa, ON, Canada) through 0.6 and subsequently 0.08 µm polycarbonate membranes (10417206 and 110604, Nuclepore Track-Etch Membrane; Whatman, GE Healthcare, Chicago, IL) sandwiched by polyester drain discs (230600, Whatman; GE Healthcare) to make size-homogenized unilamellar liposomes. We used tangential flow filtration (Krosflo KR2i; Repligen, Waltham, MA) to exchange ammonium sulfate for phosphate buffer solution (Potassium Phosphate Monobasic Anhydrous (KH_2_PO_4_; EM Science, Merck KGaA, Darmstadt, Germany) + Sodium Phosphate Dibasic Anhydrous (Na_2_HPO_4_; Sigma-Aldrich); pH 7.70) using hollow fiber filter modules (100 kDa pore size, modified polyethersulfone; Spectrum MicroKros and MidiKros, Repligen). Liposomes were syringe filtered through 0.45 µm cellulose acetate membranes (VWR). Drugs MRX-2843 (10.0 mg/mL in molecular biology grade water) and venetoclax (1.0 mg/mL in molecular biology grade water) were separately combined with liposomes and continuously mixed with magnetic stir bar at 4°C for 24 – 48 hours. Liposomes loaded with either MRX-2843 or venetoclax were then dialyzed twice in 100kDa MWCO dialysis bags (Float-A-Lyzer G2; Spectrum Laboratories, VWR) or cassettes (Slide-a-lyzer 10 MWCO G2; Thermo Scientific) in phosphate buffered saline (PBS; Corning). For long-term storage, drug-loaded lipid NPs were suspended in 99.3 mg/mL Trehalose solution (Sigma-Aldrich) in PBS. Nanoparticles were snap-frozen in liquid nitrogen and temporarily stored at -80°C. Nanoparticles were then lyophilized under low temperature (−50°C) and pressure (<0.05 mbar; Labconco Freezone, Kansas City, MO) for extended storage at -80°C. Prior to *in vitro* use, freeze-dried NPs were reconstituted in Type 1 water.

### UPLC-MS Analysis of Liposomal Drug Concentrations

Concentrations of MRX-2843, venetoclax and DSPC were measured by LC-MS using a Vanquish Horizon UPLC (Thermo Fisher Scientific) fitted with a Waters Corporation ACQUITY UPLC BEH C18 column (2.1×100 mm, 1.7 µm particle size) coupled to a high-resolution accurate mass Orbitrap ID-X Tribrid mass spectrometer. The chromatographic method for sample analysis involved elution with 20:80 ACN:water (mobile phase A) and 10:90 ACN:IPA and (mobile phase B), both in 10 mM ammonium formate with 0.1% formic acid. The following gradient program was used: 0 min 100% A; 0.5 min 100% A; 2 min 80% A; 8 min 0% A (curve 3); 10.9 min 0% A; 11 min 100% A; 12 min 100%A. The flow rate was set at 0.4 mL/min. The column temperature was set to 50 °C, and the injection volume was 2 µL. UPLC-MS^2^ experiments were performed by acquiring mass spectra with targeted MS/MS (tMS^2^) acquisition. MRX-2843 precursors, 489.3, 389.2, and 245.17 m/z, were activated with 30% HCD and analyzed in the ion trap. Venetoclax precursors 413.2, 825.4, and 868.3 m/z were activated by 35% CID and analyzed in the ion trap (40, 41). The DSPC precursor, 790.6 m/z, was selected with a 1.6 m/z isolation window, activated with 30% HCD, and the product ions were analyzed in the orbitrap at 30,000 resolution. All precursors were selected with a 1.6 m/z isolation window. Data processing was performed with Thermo Scientific Xcalibur Version 4.3.73.11. The MSMS transitions were integrated, and the data was exported to excel. The transitions for each adduct were summed and quantified from standard calibration curves.

Alternatively, drug concentrations were measured by LC-MS using an Agilent 6120 mass spectrometer (Agilent Technologies; Santa Clara, CA) with an Agilent 1220 Infinity liquid chromatography inlet. The chromatographic method involved a linear gradient of 5% acetonitrile in water (0.1% formic acid, mobile phase A) and 80% acetonitrile in water (0.1% formic acid, mobile phase B). Elution occurred over 2 minutes an Agilent Zorbax SB-C18 column (1.8 µm particle size, 2.1 mm x 50 mm) with a flow rate of 0.650 mL/min at ambient temperature. Drug concentration was quantified by integrating UV absorbance at 254 nm and normalized against an internal standard (ketoconazole) peak area.

Encapsulation efficiency (EE%) was calculated as the concentration of encapsulated drug as a fraction of the concentration of drug mixed with liposomes for pH-gradient loading. Loading capacity (wt/wt) was calculated as the mass of measured drug over the mass of the full NP (lipids + drug).

### Drug Retention post-Lyophilization

Drug retention of lipid NPs encapsulating either MRX-2843 or venetoclax following freeze-dried storage was measured by reconstituting their lyophilisates and spin filtering (100 kDa Amicon Ultracel, EMD Millipore; 100k Omega, Pall Nanosep). Drug concentrations of separate retentate and filtrate volumes were measured by UV absorbance following the Agilent 6120 LC-MS protocol described previously. Concentrations were measured in µM, where the lower limit of quantitation (LLOQ) for MRX-2843 measured at ∼ 50 µM (0.74% of measured retentate concentration), and the LLOQ for venetoclax measured at ∼ 40 µM (1.40% of retentate concentration). Drug loss is reported in µmoles.

### Lipid Nanoparticle Sizing

Liposomal size and dispersion were measured using dynamic light scattering (DLS) (DynaPro, Plate Reader III; Wyatt Technologies, Santa Barbara, CA). Prior to measurement, NPs were syringed filtered through cellulose acetate membranes (0.45 µm; VWR) and briefly bath sonicated. Particles were measured at 25°C in a black polystyrene microwell plate (Greiner Bio-One; VWR). Separately, lipid NPs were visualized with transmission electron microscopy (TEM). Using a Hitachi HT-7700 instrument at 80 kV, liposomes were added to formvar/carbon coated copper grids (400 mesh; Electron Microscopy Sciences, Hatfield, PA) in ultrapure water solution. After 15 minutes, the grids were washed with ultrapure water. Phosphotungstic acid was added to liposomes for 20 seconds for negative staining, prior to washing and imaging.

### Nanoparticle Drug Ratios

For cell-based assays, MRX-2843 and venetoclax NPs were mixed at various mol:mol ratios (M:V, 1:20, 1:5, 5:1, 20:1) Ratios were selected by comparing synergy values averaged across all combination responses within a ratio tested via HTS. Doses were filtered out for MRX-2843 (>128 nM) and venetoclax (>160 nM) that were greater than the mean AML lineage potencies (GI_50_) of 79 nM and 137 nM, respectively.

### Intracellular Drug Delivery

A mixture of monovalent MRX-2843 and venetoclax liposomes were combined at an M:V mol:mol ratio of 1:20, respectively, and incubated with OCI-AML5 cells (1×10^6^ live cells/mL in cRPMI media; 4 mL per time point) in 6-well polystyrene well plates (Greiner Bio-One; VWR). After 4 and 8 hours, cells were harvested and thrice centrifuged at 300 x g for five minutes and washed in PBS. Live and dead cell counts, including total cell mean diameters, were measured (Countess III Automated Cell Counter; Invitrogen) after the second PBS wash. Following the third round of PBS resuspension, cells were centrifuged, pelletized with aspirated supernatant, and snap-frozen over dry ice prior to storing at -80°C. In preparation for UPLC-MS analysis, frozen cell pellets were suspended in 500 µL of ice cold ACN, vortexed and sonicated (3 minutes) by cup horn (Qsonica). Samples were vacuum centrifuged for 2 hours (Savant ISS110 SpeedVac Concentrator; Thermo Fisher), reconstituted in 100 µL methanol, vortexed and sonicated by cup horn once more. For UPLC-MS, samples were mixed with 400 µL of 80% LC-MS grade methanol and 50 µL of 500 µm glass beads. Pellets were homogenized in a Tissuelyzer II (Qiagen; Maryland, USA) for 5 minutes at 30 Hz. Samples were then centrifuged at 21,100 x g for 5 minutes. Duplicate aliquots from resulting supernatants were transferred to LC vials at different dilutions for drug quantitation using the UPLC-MS protocol previously described. Intracellular drug concentrations were calculated by dividing measured drug masses by pellet volumes. Pellet volumes were calculated as the product of total cell counts (live and dead) and mean cell volumes, which were transformed from mean total cell diameters into spherical volumes.

### Apoptosis Assays

OCI-AML5 cells were cultured in 24-well plates (3×10^5^ cells/well) in cRPMI media (FBS from R&D Systems; Minneapolis, MN; Penicillin-Streptomycin from Gibco, Thermo Fisher Scientific) for 24 hours and then treated with either free drug (FD) vehicle (DMSO), nanoparticle (NP) vehicle (empty liposomes in PBS), venetoclax or MRX-2843 monotherapy, or MRX-2843 and venetoclax combined, either as FD or NP formulations for 72 hours in a total volume of 1 mL. After treatment, cells were collected, washed twice with PBS, and stained with Annexin V-FITC (5uL in 0.14 mL Annexin V Binding Buffer, #640914; Biolegend, San Diego, CA) for 10 minutes at 4°C in darkness. Propidium Iodide (#640914; Biolegend) was added (5 µL/sample) and samples were incubated for an additional 5 minutes in the dark. Annexin V (AV) and propidium iodide (PI) staining was assessed using a CytoFLEX Flow Cytometer (Beckman Coulter; Brea, CA) and FlowJo v10.8.1 analysis software (Becton Dickinson; Franklin Lakes, NJ) to detect apoptotic (AV+) and dead (AV-, PI+) cells.

### Primary AML Sample and Cell Line Screens Using Ratiometric Nanoformulations and Free Drug Combinations

De-identified bone marrow aspirates collected from pediatric patients with AML and associated non-identifying correlative clinical data were provided by the Aflac Leukemia and Lymphoma Biorepository at Children’s Healthcare of Atlanta. Samples were collected at diagnosis after written informed consent/assent and studies were conducted in accordance with relevant guidelines and regulations as approved by the Emory University Institutional Review Board (Protocol #00034535). Samples cryopreserved in freezing medium (40% RPMI 1640, 50% FBS, 10% DMSO) were thawed and washed in PBS (20% FBS v/v), then centrifuged at 600 x g (Eppendorf 5810R) for 5 minutes and suspended in medium (DMEM 85% v/v, Corning; FBS 15% v/v, VWR; 1x Penicillin-Streptomycin, VWR; 50 µM beta-mercaptoethanol, Sigma-Aldrich). 10,000 viable cells were plated per well in white polystyrene 384-well microplates (Greiner Bio-One; VWR). Cells were exposed to MRX-2843 and venetoclax (5 – 1,000 nM) alone and in combination (MRX-2843:venetoclax ratios 1:20 – 20:1) as both FD and NP formulations. Final concentrations of vehicle for FD (DMSO, 0.1% v/v) and NP (PBS, 0.1% v/v) were equal across all dosing conditions. Cell responses were further divided for analysis based on exposure to one of two cytokine maintenance medias. Cytokine media (CK) consisted of SCF (100 ng/mL), IL-6 (20 ng/mL) and TPO (10 ng/mL), while FLT-3 Rescue media consisted of CK media with added FLT-3L and IL-3 (both at 10 ng/mL), consistent with previous reports of primary AML patient sample culture (42, 43). SCF was sourced from Peprotech (#300-07; Cranbuy, NJ); cytokines IL-6, TPO and IL-3 were purchased from Prospec (IL-6, cyt-213a; TPO, enz-285-a; IL-3, cyt-210-a; Israel); and FLT-3L was sourced from Stemcell Technologies (Vancouver, BC, Canada). Once cells, drugs, and cytokines were plated, microplates were centrifuged for 5 minutes at 135 x g and then incubated for 72 hours at 37°C in a 5% CO_2_ humidified atmosphere (Electron Corporation). After 72 hours, cell viability was evaluated using the CellTiter-Glo 2.0 Assay (Promega) dispensed by the Integra Viaflo Assist Pipetting Platform (Integra Biosciences). Luminescence was measured using a SpectraMax iD3 plate reader (Molecular Devices). Viability data were background corrected to negative controls, which were empty wells containing vehicle-in-media, and normalized to positive controls, which were cells treated either with DMSO (0.1% v/v) for FD comparisons, or PBS (0.1% v/v) with the maximal lipid dose for NP comparisons. Six replicates were made for negative and positive controls, and outliers were removed by the ROUT outlier test (Q=1%). All FD and NP doses were measured in triplicate, and data was accepted for downstream calculation when Z’≥0.5 (per plate). GI_50_ is reported for logistic curve fits with R^2^>0.65. Drug synergy was calculated using the Bliss Independence model, where expected additive FD and NP effects were calculated from FD and NP monotherapies, respectively. All tested doses were drawn from the same media, FD and NP stocks, cytokine lots, and cell media prep. OCI-AML5 and MOLM-14 cells were assayed in the same manner in parallel, with the exception that cells were incubated in cRPMI media without cytokines.

### Software

Dose response curves of FD and NP monotherapies were modeled as four-parameter logistic curves with top and bottom response constraints (top = 100% GI, bottom = 0% GI) (GraphPad Prism). Growth inhibition measurements and calculations of synergy from HTS were rendered as three-dimensional surface plots in OriginPro 2022 (OriginLab Corporation; Northampton, MA). Cheminformatics analyses and molecular structures were processed on Chemicalize (Chemaxon). Experiment illustrations were drawn using BioRender. Contour plots of primary AML patient sample responses were drawn in JMP Pro, Version 16 (SAS Institute Inc.)

## RESULTS

### High-throughput Screening Identifies Synergy Between MRX-2843 and venetoclax

To maximize the therapeutic potential of combined MERTK/FLT3 and BCL-2 inhibition, we developed a formulation discovery pipeline in which pairwise drug effects were screened against leukemia cell lines in high-throughput, then utilized mathematical and computational models to (i) predict favorable drug interactions and (ii) nominate optimally potent drug pairs and mole ratios for subsequent formulation and testing against primary patient samples **(Fig 1a)**. To begin, we implemented a novel, high-throughput drug combination screen (HTS) using liquid handling robotics to construct NxN drug interaction matrices comprising the MERTK/FLT3 inhibitor, MRX-2843, and one of three BCL-2 inhibitors (BCL-2i): venetoclax, gossypol, or obatoclax. Mole ratios for each pairwise drug combination spanned 4, 5, and 6 orders of magnitude for MRX-2843:gossypol (M:G), MRX-2843:venetoclax (M:V) and MRX-2843:obatoclax (M:O), respectively.

**Figure 1.**
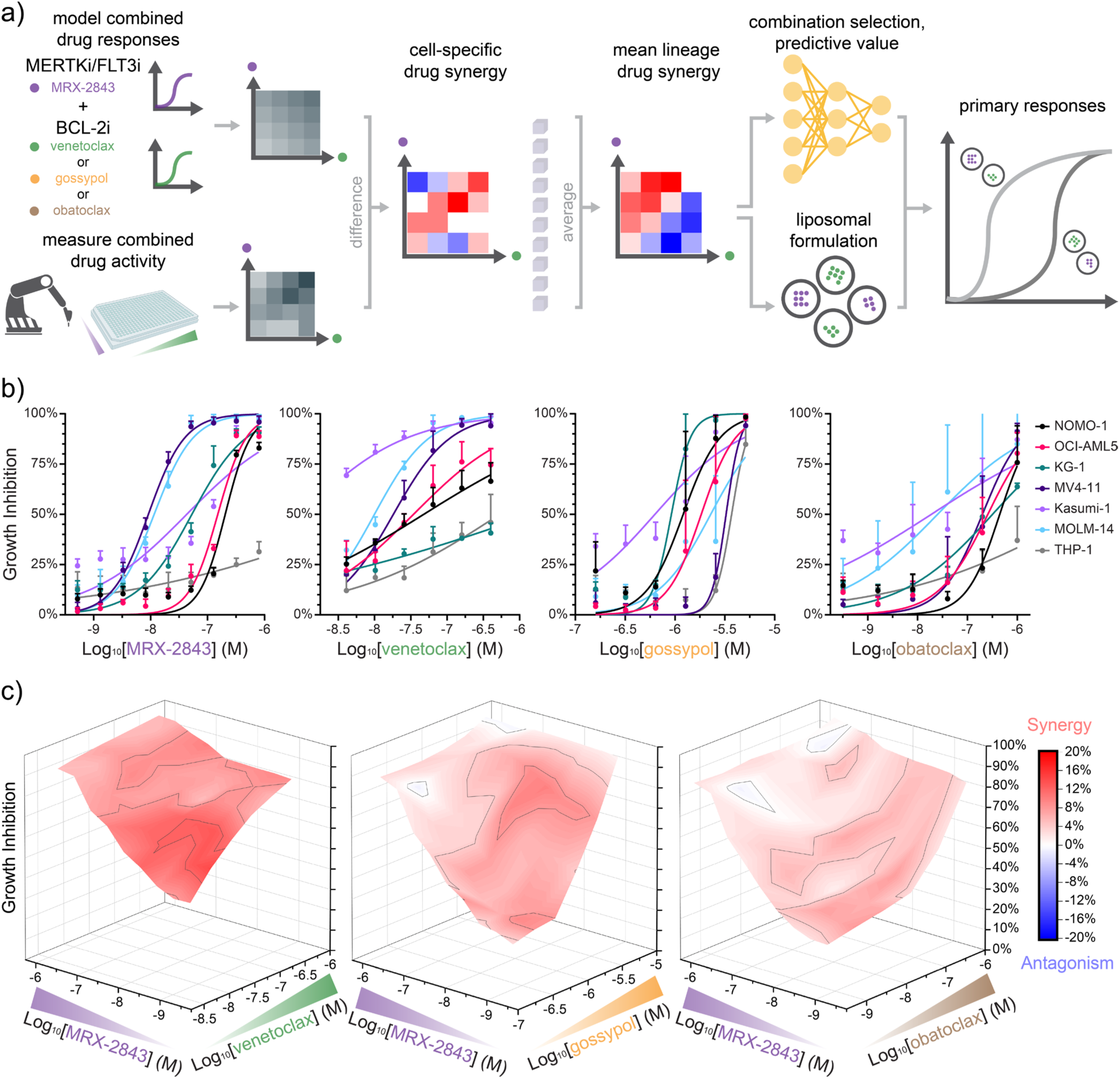
MRX-2843 and BCL-2 inhibitors synergize to reduce AML cell density in a drug ratio-dependent manner. **a)** Illustration of formulation discovery approach in which ratiometric drug synergy is screened and averaged across cell panels to inform subsequent ratiometric drug formulation and predictive pharmacogenomic correlations. **b)** Dose-dependent growth inhibition mediated by MRX-2843 and BCL-2 inhibitors, venetoclax, gossypol, or obatoclax as measured by luminescent cell viability assay (72 h, Z’≥0.5). **c)** Surface plot of drug ratio-dependent growth inhibition and synergy (color map) in AML cells treated with combined MRX-2843 and BCL-2 inhibitors (>150 discrete combinations) from screens illustrated in (a). Values in (b) represent mean ± SD of n=4 replicates. Synergy in (c) represents the percent reduction in cell density compared to the additive dose expectation modeled by Bliss Independence.

Using these drug interaction matrices, the impact of treatment on cell density was determined over 72 hours (growth inhibition, GI) in a genotypically diverse panel of AML (n=7) and infant ALL (n=3) cell lines. Cell densities were determined via bioluminescent cell viability assay with measurements collected in quadruplicate and using a Z’≥0.5 to ensure HTS data quality. Observed combination drug responses were compared to those predicted using the Bliss Independence model (44) and monotherapy drug responses measured in parallel **(Fig 1b,c S1a**,**b)**. There was strikingly consistent synergy between MRX-2843 and all BCL-2 inhibitors tested, across a range of drug ratios. The combination of MRX-2843 and venetoclax (M:V) achieved both the strongest and most synergistic responses. Hierarchical clustering of integrated synergy values indicated that the magnitude of drug synergy was cell lineage dependent with more and less responsive AML and infant ALL cell lines, respectively, clustering together **(Fig 2a)**. Further, integrated synergy responses segregated by relative specificity with pan-BCL-2 inhibitors gossypol and obatoclax clustering distinct from the BCL-2 selective inhibitor, venetoclax. Together, these data demonstrate synergistic anti-leukemia activity mediated by MRX-2843 and BCL-2 inhibitors dependent on drug ratio, cell lineage, and target specificity.

**Figure 2.**
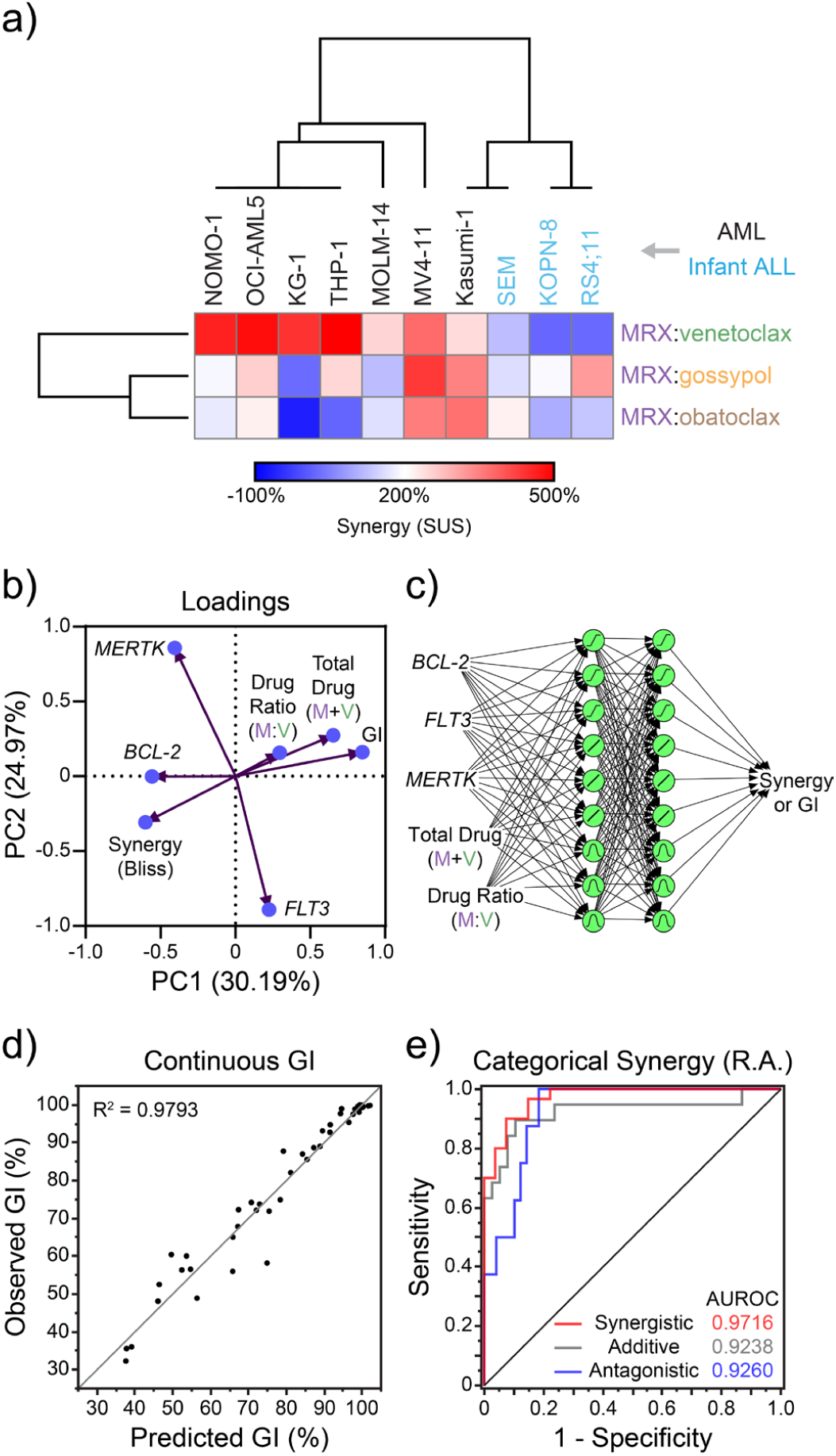
Prediction of dose-specific and ratiometric synergy between MRX-2843 and venetoclax in AML cells based on expression of genes encoding cognate drug targets and drug concentrations/ratios. **a)** Hierarchical clustering between pairwise combinations of MRX-2843 and BCL-2 inhibitors and individual cell line synergy responses (Sum of Synergy Under the Surface, SUS) **b)** Principal component analysis of variable loading vectors derived from high-throughput screening observations in AML. **c)** Schematic of two-layer perceptron used to predict synergy or GI responses based on predictor variables of MRX-2843 and venetoclax cognate gene target expression and pairwise drug combination metrics. **d)** A neural network (NN) model predicts continuous GI with high accuracy. **e)** A categorical classifier predicts synergistic, additive, and antagonistic interactions between MRX-2843 and venetoclax. (a) Cells represent the sum total of Bliss synergy, additivity, and antagonism across all pairwise combinations within a cell line (SUS). (b) Principal components (PC) are listed with the proportion of variance (%) that they account for in the HTS dataset. (b,c) Vectors and predictor variables represent log normalized gene expression (Log2(TPM+1)), total drug dose (Log2(MRX (nM) + Ven (nM)), drug ratio (Log2(MRX (nM) : Ven (nM)), combination-specifc mean measures of synergy (Bliss Independence) and GI. (d,e) Graphs are representative of 10 different dataset split vectors used to train the (d) continuous and (e) predictive classifier NN models. (e) Synergy classifications were defined as synergistic (>1% GIexp), antagonistic (<-1% GIexp), or additive by the Response Additivity (R.A.) model of drug synergy.

### Neural network-based models define the relationship between drug synergy and pharmacologic or transcriptomic features

Having shown that drug synergy was cell lineage- and therefore gene transcription-dependent, we next asked whether the relative expression of known targets of these drugs could be used to predict synergistic responses to MRX-2843 and venetoclax combination therapy *a priori*. To this end, we performed principal component analysis (PCA) on drug combination responses, incorporating the variables of drug dose, total drug (M+V), drug ratio (M:V), and target gene expression (transcripts per million, TPM) (45). The variance could best be explained by 3 principal components (PC; cumulative variance = 71.80%) **(Fig 2b, S2a)**. MERTK inversely correlated with FLT3 expression, GI clustered with increasing total drug dose (M+V) and drug ratio (M:V), and increasing GI correlated with decreasing synergy, consistent with tapering synergy as cell killing converges towards maximal GI. Interestingly, relative drug synergy clustered closely with BCL-2 expression and both metrics were inversely correlated with drug ratio and total GI. These data suggest that a mechanistic link between BCL-2 expression and drug ratio impacts the novel drug response observed here.

To quantitatively define the relationship between drug synergy and these pharmacologic or transcriptomic features, we applied machine learning to construct a nonlinear predictive model. Here, we trained two-layer neural network (NN) models using 5 predictor variables to predict Synergy or GI **(Fig 2c)**. Prediction of total GI was highly accurate (R^2^=0.9793), as was the binary classification of synergy (AUROC>0.99) **(Fig 2d, S2b)**. We challenged the predictive classifier by replacing the Bliss model of drug synergy with Response Additivity (46), which added an antagonistic class to synergy responses. Even with this model exchange, predictions were consistently high **(Fig 2e)**, with AUROC>0.92 across all three synergy classes (antagonistic, additive, or synergistic). While continuous synergy prediction was also possible, the NN models predicted continuous synergy values less accurately (R^2^=0.6828 for response additivity; R^2^=0.7658 for Bliss) than categorical synergy **(Fig S2c-e)**. Statistical simulations indexed the relative importance of predictor variables on output accuracy, showing that total drug (M+V), target gene expression (BCL-2, MERTK, FLT3), and drug ratio (M:V) were significant in the predictive classification of synergy **(Fig S2f)**.

### Combination formulations of MRX-2843 and venetoclax recapitulate the pharmacology and synergy observed in high-throughput screens

We next devised a drug formulation strategy to maximize the therapeutic potential of combined MRX-2843 and venetoclax via delivery of a fixed, synergistic ratio. Using *in silico* models of pH-dependent drug charge and lipophilicity (logP) **(Fig S3a)**, we developed a single pH gradient-based process by which either drug could be remote-loaded within lipid vesicles **(Fig 3a)** comprising DSPC:DSPG:cholesterol (7:2:1 molar ratio), a lipid mixture commonly utilized in clinically approved lipid drug formulations (47). Following purification, MRX-2843 or venetoclax formulations were lyophilized in trehalose and later reconstituted in deionized water with >97% drug retention as measured by LC-MS **(Fig S3b)**. Formulations demonstrated high batch-to-batch encapsulation reproducibility **(Fig 3c)** and maintenance of hydrodynamic size and polydispersity as measured by DLS **(Fig 3b)**.

**Figure 3.**
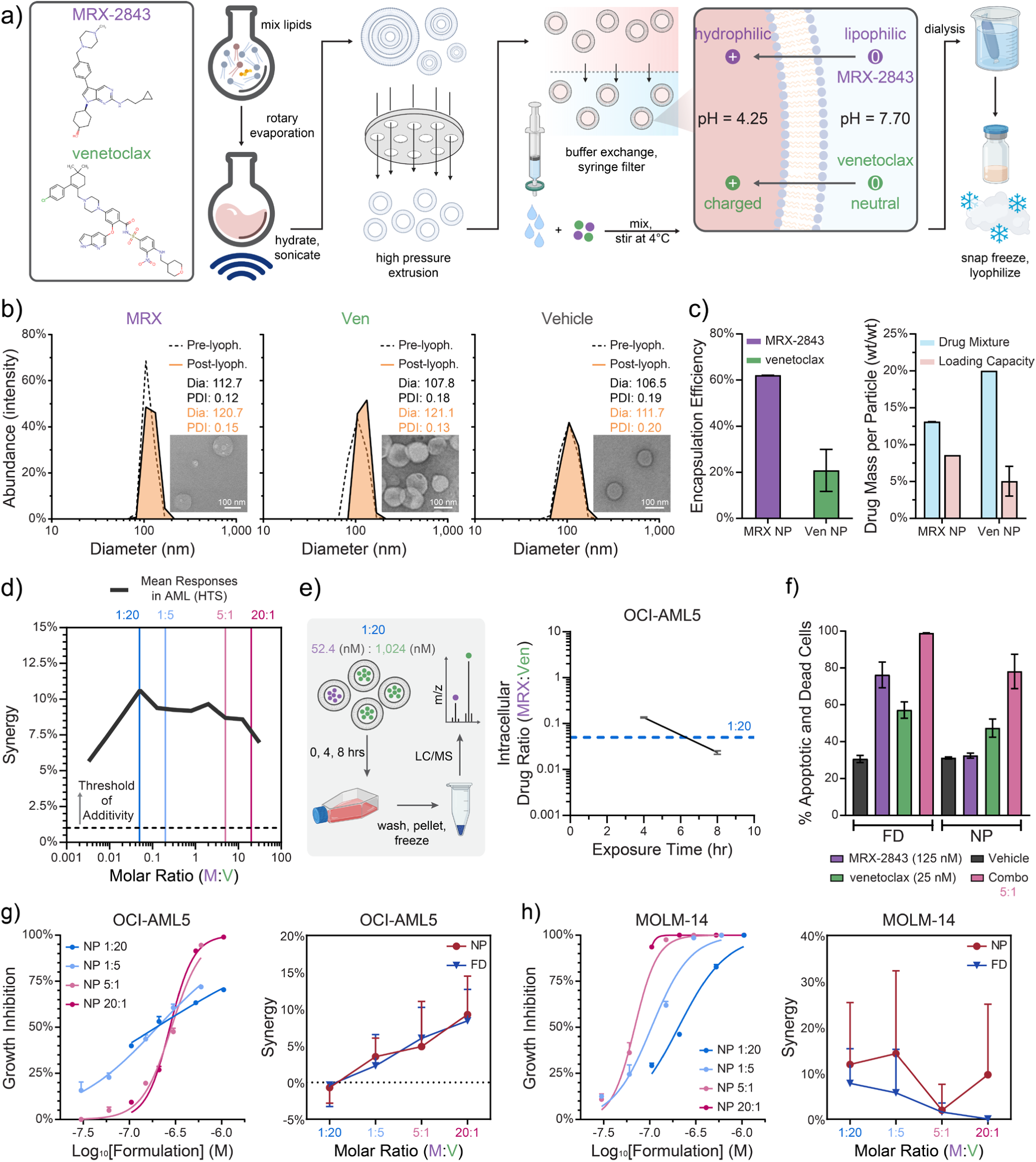
Combination formulations of MRX-2843 and venetoclax nanoparticles maintain drug ratios and associated synergy following cell delivery. **a)** illustration of MRX-2843 and venetoclax nanoparticle formulation via remote-loading into unilamellar lipid vesicles. **b)** Hydrodynamic size distribution and polydispersity index before and after freeze dried storage as measured by dynamic light scattering. Insets illustrate pre- and post-lyophilization morphology as imaged by transmission electron microscopy. **c)** Drug encapsulation efficiency and percent loaded by weight measured by LC-MS. **d)** Plot of mean drug synergy (Bliss model) as a function of mole ratio (MRX-2843:venetoclax) from the data in Fig 1. Synergistic mole ratios of 1:20, 1:5, 5:1 and 20:1 (vertical lines) were selected for subsequent assays. **e)** Kinetics of drug delivery in OCI-AML5 cells following treatment with a 1:20 (MRX-2843:venetoclax) nanoparticle formulation measured by LC-MS. **f)** Induction of apoptosis and cell death in OCI-AML5 cells treated for 72 hrs with single agents or a 5:1 molar ratio of MRX-2843 and venetoclax combined as either free drug (FD) or nanoparticle (NP) formulations measured by flow cytometry after staining with Annexin V and propidium iodide. **g**,**h)** Dose-dependent growth inhibition (left panels) and mole ratio-dependent synergy (right panels) mediated by combination nanoparticle formulations in **(g)** OCI-AML5 and **(h)** MOLM-14 cells demonstrating maintainence of drug potency and ratiometric synergy following nanoparticle formulation and storage. (c) Mean values ± SD from two nanoparticle batches. (e) Mean values ± SD of two technical replicates. (f) Mean values ± SEM of single replicates from two independent experiments. (g,h) Mean ± SD of triplicate measures for growth inhibition and mean ± SEM of n=4-5 tested doses within each indicated mole ratio.

To determine whether ratiometric mixtures of MRX-2843 and venetoclax NPs recapitulated the synergistic effects of their constitutive free drug components, four ratiometric combinations were selected for *in vitro* testing (1:20, 1:5, 5:1, 20:1 M:V) **(Fig 3d)**. Strikingly, a 1:20 M:V ratio achieved cytosolic levels of MRX-2843 and venetoclax in OCI-AML5 cells that closely mirrored those in the nanoparticles (exp: 0.05, obs: 0.022 – 0.138 M:V) **(Fig 3e, S3c)**. Both FD and NP formulations delivering a 5:1 ratio of MRX-2843 and venetoclax induced apoptosis in OCI-AML5 cells as measured by annexin v staining and both FD and NP combinations were more effective than their respective single agent components **(Fig 3f, S3d)**. Finally, we systematically compared dose-dependent GI mediated by NP and FD formulations at different ratios in OCI-AML5 (FLT3 wild-type) and MOLM-14 (FLT3-ITD+) AML cells. While single-agent FDs were nominally more potent than single-agent NPs over the concentration gradients tested **(Fig S3e)**, both formulations of pairwise combinations mediated increased efficacy with increasing M:V ratios **(Fig 3g,h**, **S3f)**. Synergy was also ratio-dependent in both OCI-AML5 and MOLM-14 cells, with strong concordance between FD and NP formulations. Thus, combination nanoformulations of MRX-2843 and venetoclax constitutively maintain ratiometric drug synergy following *in vitro* cell delivery.

### Ratiometric Nanoformulations are Constitutively Synergistic in Primary AML Patient Samples

To assess the translational potential of MRX-2843 and venetoclax combination nanoformulations, the impact of treatment with four different molar ratios of NPs was determined in primary AML patient samples *ex vivo* **(Fig 4a)**. Cryopreserved, diagnostic bone marrow specimens from 5 different pediatric patients with AML and different prognostic (e.g. high or low risk, FLT3-ITD+) or clinical features (e.g. subsequent relapse, central nervous system involvement) **(Fig S4a)** were treated with FD or NP combinations and compared to equivalent single-agents. Two different media preparations (with and without FLT3 ligand) were also used to mimic either basal or mitogenic growth conditions.

**Figure 4.**
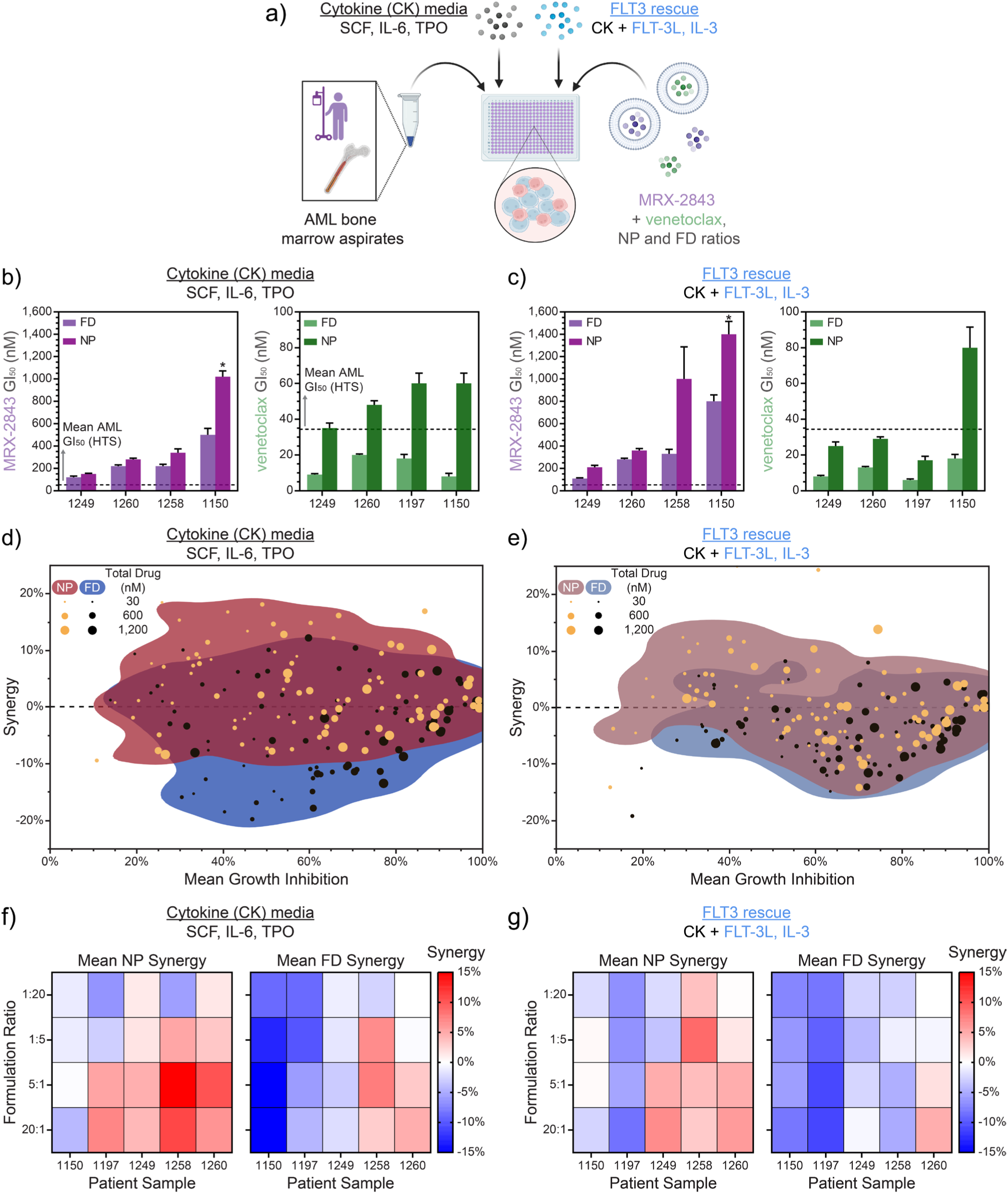
Ratiometric nanoformulations of MRX-2843 and venetoclax mediate synergistic anti-leukemia activity in primary AML patient bone marrow samples. **a)** Schematic of approach to test nanoparticle (NP) and free drug (FD) MRX-2843 and venetoclax formulations in primary AML patient samples. **b**,**c)** Dose-dependent growth inhibition mediated by FD and NP formulations of single-agent MRX-2843 or venetoclax demonstrating maintenance of drug potency following NP formulation as measured by luminescent cell viability assay (72, Z’≥0.5) in **(b)** basal or **(c)** FLT-3L supplemented media. **d**,**e)** Contour plot of NP (red/brown) and FD (blue) growth inhibition versus synergy in primary AML patient samples as measured by luminescent cell viability assay (72 h) in **(d)** basal or **e)** FLT3L-supplemented media. Individual FD and NP doses are represented by yellow and black dots, respectively. **f**,**g)** Heatmaps showing synergy as a function of M:V ratio across AML patient samples cultured in **(f)** basal or **(g)** FLT-3L supplemented media. Values in (b,c) represent mean GI50 +/-SEM of three technical replicates. Hash lines in (b,c) show mean GI50 values from Fig 1b and asterisks denote extrapolated GI50 values. (f,g) Color map shows synergy averaged across all doses for a given ratio, per patient.

Consistent with results in AML cell lines, both MRX-2843 and venetoclax NPs had potent, dose-dependent anti-leukemia activity in primary AML cell cultures **(Fig 4b,c**, **S4b-d)**. *Strikingly*, ratiometric combination formulations were synergistic in four of five patient samples, whereas corresponding free drug combinations were predominantly antagonistic **(Fig 4d-g)**. Of note, these differences were less pronounced in cell growth media supplemented with FLT-3 ligand and FLT3 is an MRX-2843 target. The mean synergy of both FD and NP formulations was ratio-dependent **(Fig 4f,g)** with 5:1 and 20:1 M:V ratios providing superior synergy compared to lower ratiometric doses and NP formulations providing more favorable interactions relative to the FD combinations across all patients **(Fig S4e)**.

## DISCUSSION

Here we address the unmet clinical need to develop new, more effective combination therapies for AML. Treatment with the dual MERTK and FLT3 inhibitor, MRX-2843, synergized with pharmacologic inhibition of anti-apoptotic BCL-2 family proteins to decrease AML cell expansion *in vitro*. Systematic, high-throughput combination drug screens revealed that MRX-2483 most profoundly and consistently synergizes with the BCL-2 specific inhibitor, venetoclax, and to a lesser extent the pan BCL-2 inhibitors gossypol or obatoclax (48, 49) to provide anti-leukemia activity across a genotypically diverse panel of AML cell lines and primary patient samples. These findings are consistent with recent reports demonstrating synergistic interactions between venetoclax and FLT3, dual FLT3/AXL, or dual MERTK/AXL inhibitors in AML (50-52) and provide the first demonstration of synergy between a dual MERTK/FLT3 inhibitor and venetoclax in AML. Of note, MRX-2843 synergized with venetoclax in the absence of activating FLT3 mutations. These data provide the first evidence implicating BCL-2 as a mediator of resistance to MERTK inhibition in AML and implicate MERTK inhibition as a mechanism of synergy with venetoclax. Moreover, we demonstrate that combined drug synergy is strongly ratio-dependent, a finding that is significant given that plasma ratios of co-administered small molecule drugs often fluctuate dramatically, even when administered concurrently.

To exploit and maximize the therapeutic benefit of combined MRX-2843 and venetoclax delivery, we developed nanoscale drug formulations encapsulated within lipid vesicles that maintain synergistic drug ratios and deliver them intracellularly. The composition of these particles closely mirrors that found in FDA-approved nanomedicines, including DaunoXome, Lipoplatin, Vyxeos and Onivyde (53, 54), with the goal to facilitate clinical translation. Of note, this approach may harmonize and augment delivery of MRX-2843 and venetoclax *in vivo* as prior studies of liposomal Vyxeos demonstrated preferential delivery of its nanoparticle contents, cytarabine and daunorubicin, to the bone marrow, a common reservoir of disease in AML (15, 55, 56). The remote, pH gradient-based loading is also significant in that this approach is intrinsically scalable, as demonstrated by the commercial scale production of DaunoXome and Doxil (57). The ability to lyophilize and reconstitute liposomal MRX-2843/venetoclax with minimal drug loss (<3% total drug) or shift in product attributes further demonstrates strong translational and commercial potential.

MRX-2843/venetoclax nanoformulations also had potent therapeutic activity against primary AML patient samples, indicating their potential for translational application. Moreover, both the magnitude and frequency of synergistic responses were improved in patient bone marrow samples treated with nanoformulations compared to treatment with free drugs. Further, nanoformulations provided superior synergy compared to free drugs under culture conditions associated with FLT3 inhibitor resistance (i.e. FLT3L stimulation (58)). These data implicate nanoparticle delivery of ratiometric MRX-2843 and venetoclax combination therapy as an effective strategy to treat AML with or without activating FLT3 mutations, with potential to prevent or overcome resistance to FLT3 inhibitor monotherapies.

Results from the high throughput combination drug screen were also combined with cell-specific transcriptomic profiles to develop a classifier that predicts synergy between MRX-2843 and venetoclax based on total and ratiometric drug exposure, as well as MERTK, FLT3, and BCL-2 mRNA expression levels. Co-development of predictive biomarkers will be critical to identify patients who may benefit from MRX-2843 and venetoclax combination therapy. Personalized disease models based on transcriptomic and drug sensitivity profiling have already been used to predict response to venetoclax in two pediatric leukemia patients (59). While expansion of training datasets for the model described here and its prospective evaluation are in progress, this approach may find utility in the future as a companion diagnostic to guide patient or trial participant selection.

## CONCLUSION

In summary, we demonstrate a systematic approach to combination drug screening and nanoscale formulation that maximizes drug synergy between the small molecule MERTK/FLT3 inhibitor MRX-2843 and BCL-2 family protein inhibitors. MRX-2843 synergized with venetoclax in high-throughput drug screens to inhibit AML cell expansion in a ratio-dependent manner. To exploit these findings, we developed lipid NP formulations of MRX-2843 and venetoclax that maintain optimal drug stoichiometry and deliver ratiometric doses in AML cells. Nanoscale formulations mediated potent anti-leukemia activity and provided enhanced synergy compared to equivalent FD combinations in primary AML patient samples, indicating their potential as therapeutic agents. They also have favorable properties for clinical application, including stable storage of lyophilized product, a lipid nanoparticle formulation that is FDA-approved for patients with AML, and co-development of a pharmacogenomic model to predict synergistic response. With further research, these findings may lead to development of new, personalized treatment options for patients with relapsed or refractory AML.

## ACKNOWLEDGEMENTS

We are grateful for assistance from the Pediatric General Equipment Core, the Robert P. Apkarian Integrated Electron Microscopy Core, the Emory University Lipidomics Core, the Emory University School of Medicine Flow Cytometry Core, the Georgia Institute of Technology Systems Mass Spectrometry Core Facility, and the Emory Chemical Biology Discovery Center. Patient samples were provided by the Aflac Leukemia and Lymphoma Biorepository at Children’s Healthcare of Atlanta; other investigators may have received specimens from the same subjects. The content here is solely the responsibility of the authors and does not necessarily represent the official views of the organizations acknowledged herein.

## AUTHOR CONTRIBUTIONS

J.M.K., J.J., A.T., R.J.S., X.W., N.T.J., H.F., Y.D., D.D., D.K.G., and E.C.D. designed research; J.M.K, J.J., A.T., M.Q., L.A.B., S.G.M., H.Z., E.C., and B.U. performed research or analyzed data; J.M.K., J.J., Y.D., D.D., D.K.G., and E.C.D. wrote or edited the manuscript.

