## Supplementary Information for "Constitutively synergistic multiagent drug formulations targeting MERTK, FLT3, and BCL-2 for treatment of AML"

### This file includes:

Fig. S1: *MLL*-rearranged (infant) ALL Ratiometric Screening Responses
Fig. S2: Computational Predictions of MRX-2843 and venetoclax Responses
Fig. S3: Characteristics of Nanoformulations
Fig. S4: Free Drug and Nanoformulation Responses in Primary AML Patient Samples

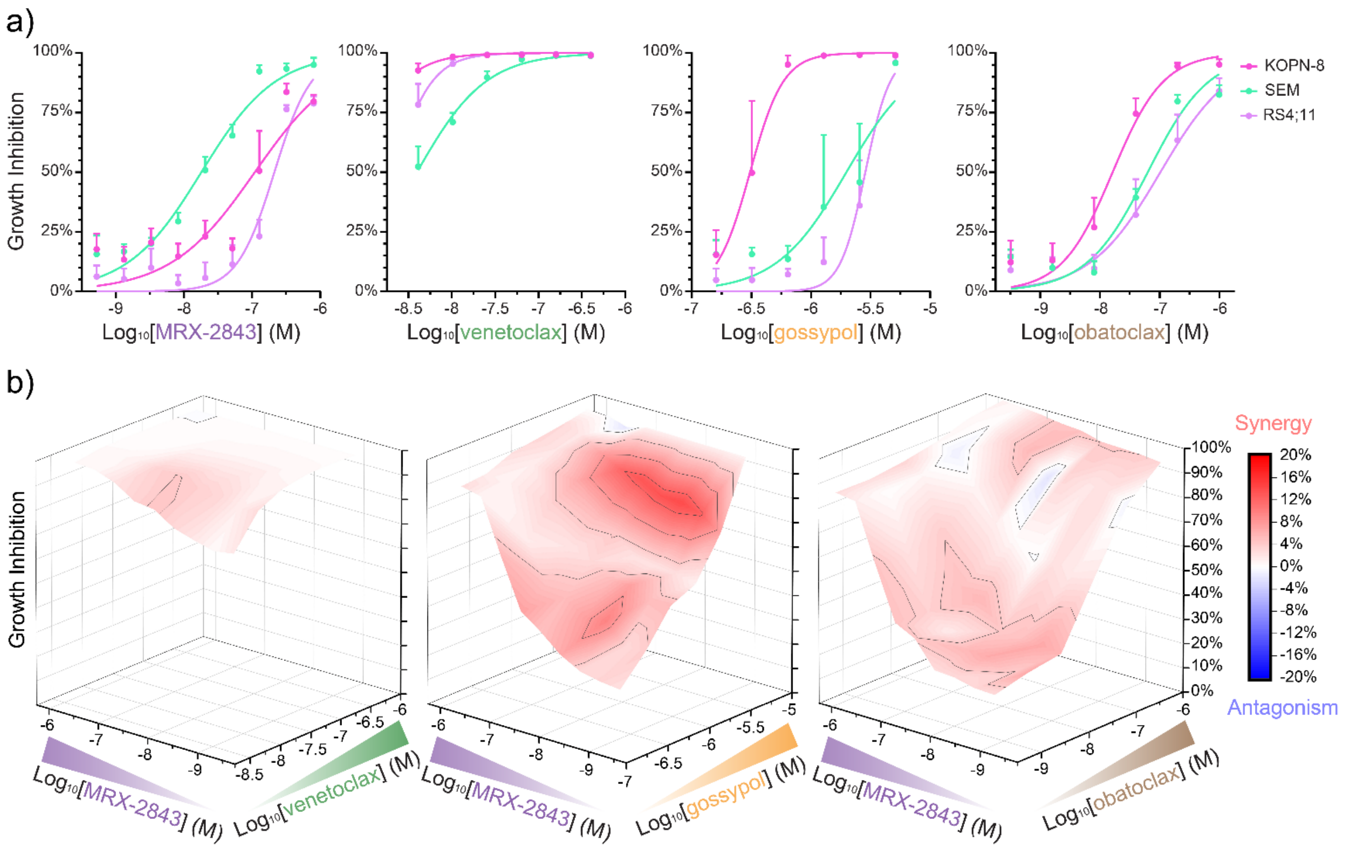


**Fig. S1. MRX-2843 synergizes with BCL-2 inhibitors in *MLL*-rearranged B-ALL. a)** Single-agent MRX-2843 and BCL-2 inhibitor responses are shown in Infant ALL cell lines as measured by luminescent cell viability assay (72 h, Z'≥0.5) **b)** Mean growth inhibition and pairwise combination synergy between MRX-2843 and BCL-2 inhibitors are shown. (a) Mean values ± SD of n=4 replicates are shown. (b) Surfaces trace the mean inhibition of cell expansion across all MLL-rearranged precursor B-ALL cell lines for >150 distinct drug combinations. Synergy (red) represents the percent reduction in cell density compared to the additive dose modeled by Bliss Independence.

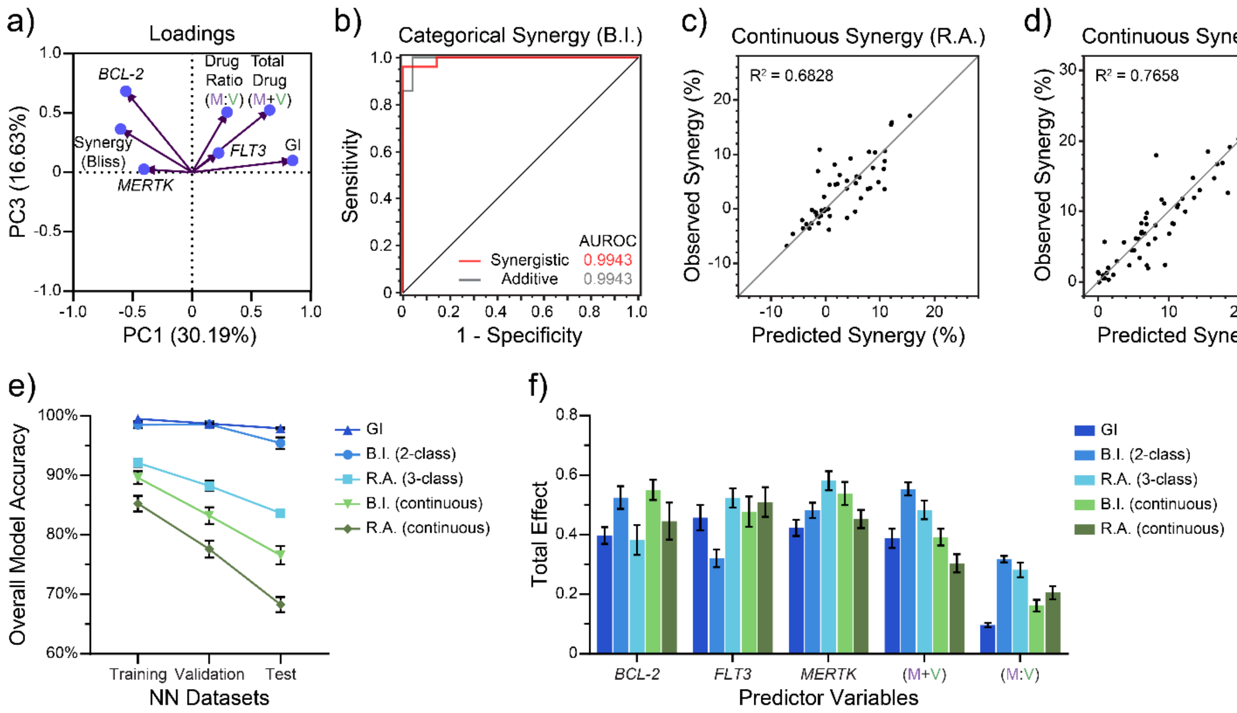


**Fig. S2. Predictor variables comprising target gene expression and metrics of MRX-2843 and venetoclax drug combinations allow for continuous and categorical prediction of pairwise drug synergy in neural network models. a)** Principal component analysis depicting principal components (PC) PC1 and PC3 with cumulative proportion of variance 71.80%. **b)** Neural network (NN) two-class predictive classifer of drug synergy using the Bliss Independence (B.I.) model. **c)** Continuous predictions of synergy using the Response Additivity (R.A.) and **d)** B.I. models. **e)** Accuracy comparisons of continuous NN models and categorical classifiers with respect to Training, Validation and Test sets (70:15:15 split). **f)** Predictor variables are scored based on their relative importance in affecting output predictions of NN models. (b-d) Graphs are representative of 10 different dataset split vectors used to train the (c,d) continuous and (b) predictive classifier NN models. (e,f) Data represent mean values ± SEM of n=10 independent tests evaluated with distinct partitions of the AML cell line high-throughput screening data into Training, Validation and Test sets. (f) Total effects quantify relative importance of predictor variables alone and when grouped with others as assessed by Monte Carlo simulations of independently resampled inputs (JMP).

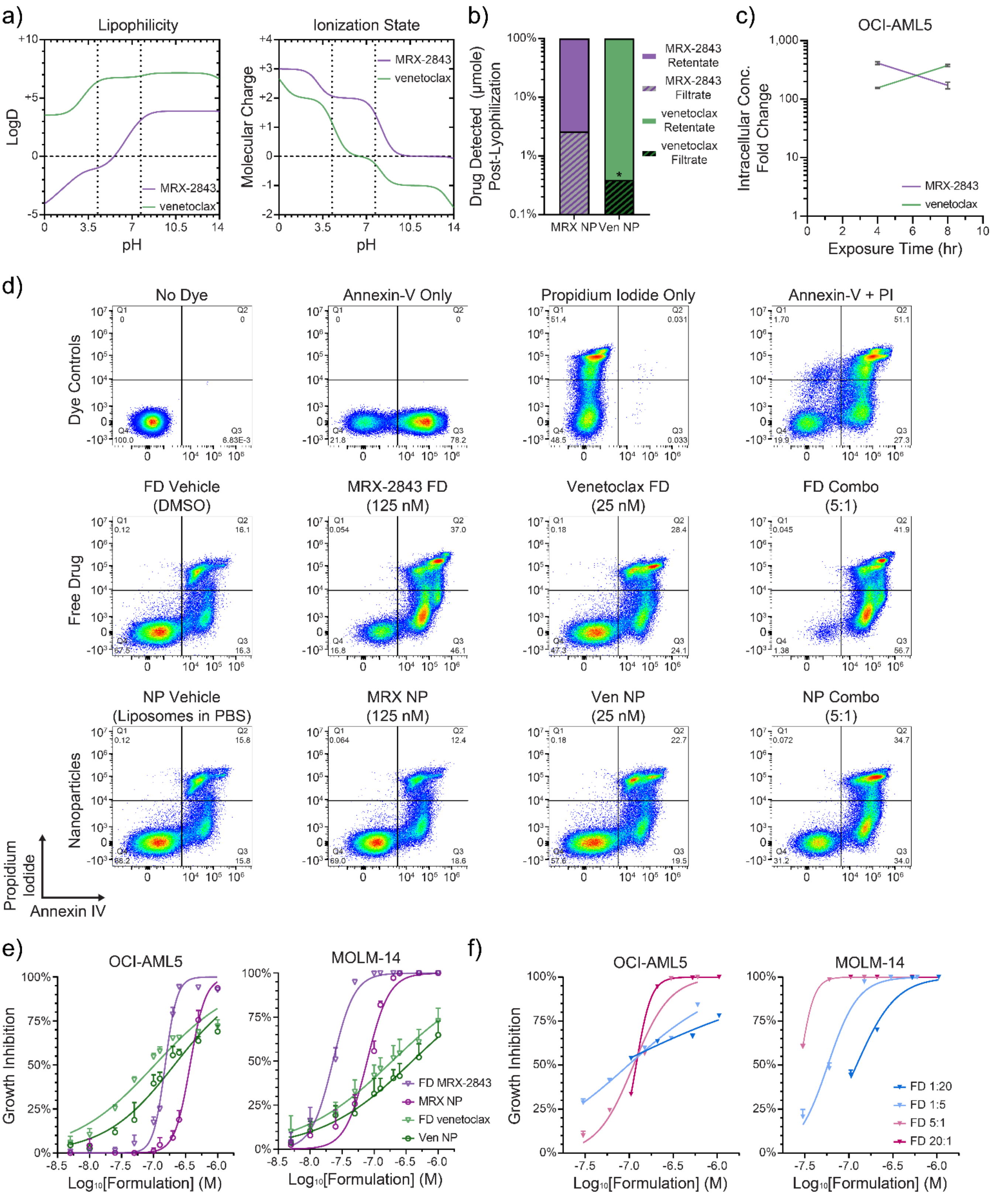


**Fig. S3. MRX-2843 and venetoclax nanoparticles exhibit high drug retention, efficient drug delivery, and achieve similar induction of apoptosis and growth inhibition in AML cell lines compared to free drug formulations. a)** Dashed lines at pH = 4.25 and pH = 7.70 interpolate lipophilicity and molecular charge of MRX-2843 and venetoclax that may be exploited for pH-gradient drug loading into liposomes as predicted by cheminformatics. **b)** Following reconstitution of MRX-2843 and venetoclax nanoparticle lyophilisates, drug loss was measured by LC-MS. **c)** Intracellular MRX-2843 and venetoclax concentrations in OCI-AML5 cells are shown as fold change relative to initial NP concentrations in solution as measured by LC-MS. **d)** Representative flow cytometry profiles showing cell death (Q1+Q2+Q3) after treatment with either free drug (FD) or nanoparticle (NP) formulations of 5:1 MRX-2843:venetoclax as measured by flow cytometry. **e)** FD and NP single-agent responses are shown in OCI-AML5 and MOLM-14 cell lines as measured by luminescent cell viability assay (72 hr). **f)** Ratio-dependent differences in FD formulations are shown in OCI-AML5 and MOLM-14 cell lines. (a) Data is reproduced by cheminformatics analysis (Chemicalize.com). (b) Data is measured from a single nanoparticle batch, where * denotes estimation of venetoclax Filtrate occurring below the lower limit of quantitation (LLOQ). (c) Error bars show the range of fold change in intracellular drug concentrations compared to the mixing concentration from duplicate samples at 4 and 8 hours. (d) Data shows representative quadrant event densities used to calculate apoptotic cells via flow cytometry. (e,f) Mean values ± SD of (e) three replicate dose responses or (f) n=4-5 different pairwise concentration combinations per ratio.

a)

| Pt. Sample | PB Blast (%) | BM Blast (%) | MRD (%), EOI | Risk | Relapse | Second Relapse | CNS Disease | Normal Cytogenetics | Molecular and Prognostic Markers |
|---|---|---|---|---|---|---|---|---|---|
| 1150 | 87.3 | 75.8 | 0.7 | HR | Yes | No | No | No | --- |
| 1197 | 95.7 | 96.4 | Negative | LR | No | No | Yes | Yes | Nucleophosmin |
| 1249 | 94.7 | 90.6 | 39 | HR | Yes | Yes | No | No | FLT3-ITD, Monosomy 7 |
| 1258 | 85.6 | 81 | Negative | LR | No | No | Yes | No | --- |
| 1260 | 90.5 | 96.4 | 86.6 | HR | Yes | Yes | No | No | *MLL*-rearrangement |

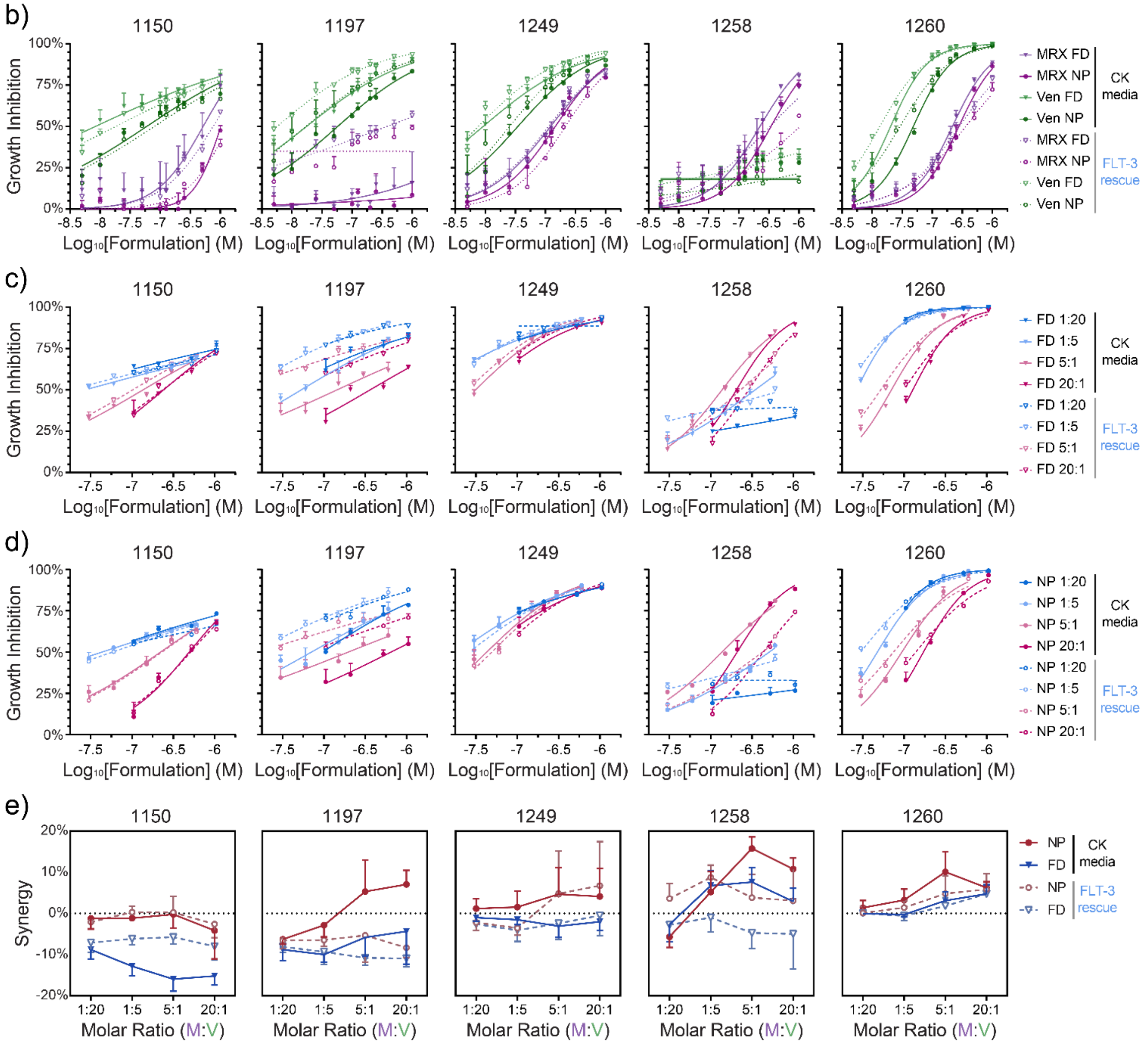


**Fig. S4. Nanoparticle formulations of combined MRX-2843 and venetoclax provide dose-dependent anti-leukemia activity and enhanced ratio-dependent synergy compared to free drug. a)** Correlative clinical data associated with five independent AML patient samples, including peripheral blood (PB) and bone marrow (BM) blast percentage at diagnosis, minimal residual disease (MRD) at end of induction (EOI), high risk (HR) or low risk (LR) status, and presence of disease in the central nervous system (CNS) at diagnosis. **b)** Single-agent dose responses in primary AML patient samples treated with free drug (FD) or nanoparticle (NP) formulations as measured by luminescent cell viability assay (72 hr). **c,d)** Growth inhibition as a function of MRX-2843:venetoclax ratio in AML patient samples treated with **(c)** FD or **d)** NP combination formulations. **e)** Mean synergy as a function of MRX-2843:venetoclax ratio in AML patient samples treated with FD or NP combination formulations. (b-d) Mean values ± SD of triplicate dose measurements are shown. (d) Data represent mean synergy values ± SD averaged across all dose responses within a ratio.
